# A compact protein panel for organ-specific age and chronic disease prediction

**DOI:** 10.1101/2024.12.09.627624

**Authors:** Anastasiya Vladimirova, Ludger J.E. Goeminne, Alexander Tyshkovskiy, Vadim N. Gladyshev

**Author notes:** Correspondence should be addressed to V.N.G.

## Abstract

Recent advances in plasma proteomics have led to a surge of computational models that accurately predict chronological age, mortality, and diseases from a simple blood draw. We leverage the data of ∼50,000 participants in the UK Biobank to investigate the predictive power of such models compared to individual proteins and metabolites by assessing disease risk and organ aging. We find that, with the exception of brain-related diseases, individual protein levels often match or surpass the predictive power of elaborate clocks trained on chronological age or mortality risk. Certain proteins effectively predict multiple diseases affecting specific organs. We show that in most cases, proteins predict diseases better than polygenic risk scores, and identify novel associations between human plasma protein levels and diseases, including LAMP3 and COPD, CHHR2 and liver disease, FAMC3 and kidney disease, and TMED1 and gout. We present a focused panel of 21 protein biomarkers that reveals the health state of the six organs associated with major age-related diseases. Our panel predicts common age-related diseases, including liver cirrhosis and fibrosis, dementia, kidney failure, and type II diabetes better than established blood panels and aging models. Through its vast coverage of age-related diseases, our compact panel offers a cost-effective alternative to full-scale proteomic analyses, making it a prime candidate for the non-invasive clinical detection and management of numerous age-related diseases simultaneously.

## Introduction

The relationship between aging and diseases is complex and involves multiple physiological processes. While the precise mechanisms linking aging and disease development remain an area of active investigation, with several theoretical frameworks proposed, aging manifests or leads to frailty, functional decline, and increased likelihood of diseases, such as cancer, cardiovascular disease, neurodegenerative disorders, and metabolic diseases (Cohen et al., 2020; Gladyshev, 2016; Hodes et al., 2016; Li et al., 2023).

Quantifying protein concentrations in blood plasma provides a non-invasive means for evaluating systemic health(Sun et al., 2023) . Since blood circulates throughout the body and includes components derived from various organs (Jiang et al., 2020), it reflects the physiological and pathological status of various organs, thereby providing valuable insights into disease states and organ dysfunction. Pioneering work by Oh et al., 2023 used innovative blood proteomic panels to show that organs may age at different rates and that accelerated organ aging is linked to organ-specific diseases. Argentieri et al., 2024 then utilized the Olink Panel from ∼50 000 participants that was recently provided by UK Biobank (Sun et al., 2023) to train proteomic aging clocks on all 2,897 Olink proteins in the panel, while we and others (Goeminne et al., 2024; Oh et al., 2024) extended this approach by applying organ-specific aging models to the UK Biobank dataset, training them on mortality and revealing numerous associations with lifestyle, environment and disease. Hence, we proposed that chronic diseases are representations of accelerated organ-specific aging (Goeminne et al., 2024).

Various plasma proteins have been established as robust disease biomarkers (Berkowitz, 2004; Wan C Fu, 2024; Winchester et al., 2023), and others have already extensively characterized the predictive power of such proteins, alone or in combination, for a variety of diseases reported in the UK Biobank (Carrasco-Zanini et al., 2024; Gadd et al., 2024; You et al., 2023). While You et al., 2023 and Carrasco-Zanini et al., 2024 utilize machine learning algorithms, like LASSO and neural networks, to leverage the most information in predicting disease risk, Gadd et al., 2024 characterizes each protein’s individual risk prediction capabilities. Furthermore, Gadd et al., 2024; Julkunen et al., 2023 use the UK Biobank nuclear magnetic resonance (NMR) metabolic biomarker data to describe the associations between metabolites and diseases for more than 100,000 patients. Despite advancements in the development of disease biomarkers, a direct comparison of the predictive performance of aging clocks versus specific protein and metabolite panels has not been previously reported to the best of our knowledge. A concise panel of molecular biomarkers encompassing the majority of common age-related diseases and overall mortality has also been lacking.

In this study, we conduct a comprehensive analysis of the predictive power of individual proteins, metabolites, and models based on different subsets of proteins trained to predict both chronological aging and mortality, as well as models trained to directly predict diseases occurrence risk. With the exception of brain diseases like dementia, we show that single proteins perform either *on par* or better at predicting disease risks compared to state-of-the-art plasma-protein-based aging models, both conventional, which are trained on all available protein features, and organ-specific.

We present a concise panel of twenty-one protein biomarkers that assesses organ-specific aging of the six organs (brain, heart, lung, liver, kidney, and pancreas) as well as general mortality, and accurately predicts common age-related diseases. The seven protein biomarkers selected for brain aging are features of the small feed-forward neural network trained to predict chronological aging. This brain panel demonstrates significant improvement in predicting brain aging and its associated pathologies, both over single protein biomarkers and more elaborate models. The proteins in our panel remain stable predictors, even after adjusting for environmental and other factors (Garg et al., 2023). With its broad coverage, our panel holds a tremendous clinical potential for the low-cost early detection of a plethora of age-related diseases simultaneously.

## Results

### Single proteins can outperform organ-specific aging models in predicting organ-specific diseases

To evaluate the predictive performance of organ-specific models and individual protein levels for disease risk, we leveraged proteomics data from 44,952 UK Biobank participants with over a decade of post-sample health data collection (Fig. 1A). By comparing normalized residuals of predicted age/mortality risk scores with normalized single protein levels, that are also adjusted for age, we assessed their ability to predict mortality and 10 major age-associated diseases (heart failure, kidney failure, chronic obstructive pulmonary disease (COPD), liver cirrhosis, stroke, myocardial infarction, hypertension, dementia, type II diabetes, and cancer), as well as an extended list of 373 diseases available from UK Biobank.

**Figure 1.**
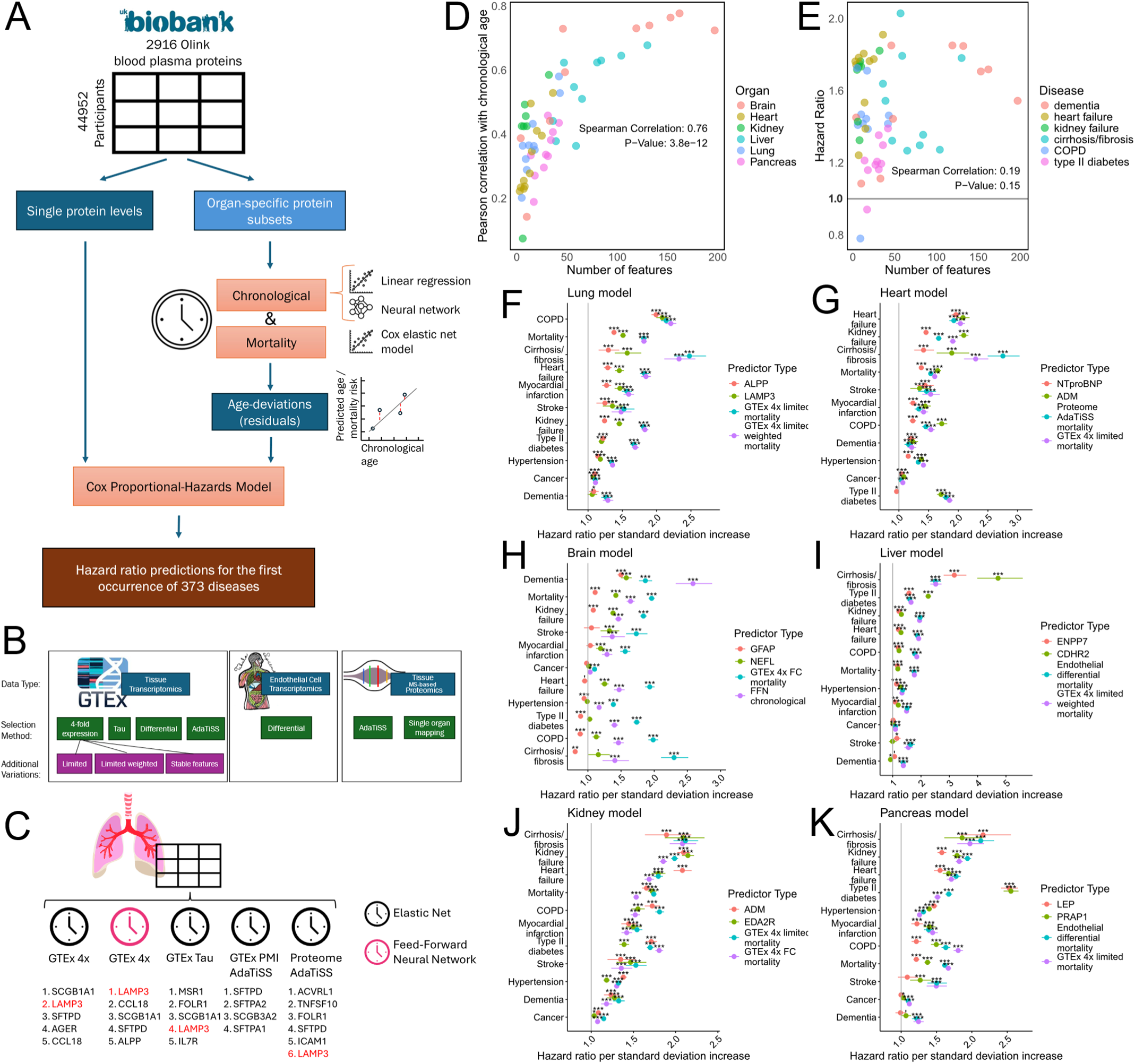
Single-protein predictors generally outperform organ-specific aging models in predicting organ-specific diseases. A. Overview of the analysis pipeline. The UK Biobank proteomics dataset (n = 44,952 participants) is used to predict chronological age and mortality. The models’ age deviations and single protein levels are compared in their ability to predict 373 common diseases. **B**. Overview of approaches for selecting proteins for organ-specific aging models. These include three different types of data: trasncriptomics (GTEx Consortium, 2020), single-cell transcriptomics (Consortium* et al., 2022) and proteomics (Jiang et al., 2020; Prakash et al., 2022), 5 different computational methods for organ specificity, and three additional variations of the GTEx 4x approach, see Methods. **C**. Proteins with the largest absolute contribution (coefficients multiplied by average protein levels) to lung-specific chronological models. Colors represent the type of the computational model (elastic net vs feed-forward neural network). LAMP3 (red) overlaps across nearly all models, if it’s present in the feature set. **D-E**. The horizontal axis shows the number of proteins included in di@erent types of models (as described in panel **B**) for six organs and these models’ predictions’. The vertical axis shows the model predictions’ correlation with age (**D**) or the hazard ratios for a common age-related disease a@ecting the organ in question (**E**). The grey line indicates a hazard ratio of 1. **F-K.** Best significant organ-specific models and best significant single protein predictors, sorted by hazard ratios, of 10 comon age-related diseases and mortality in selected organs. The hazard ratios for age-related diseases per standard deviation increase in the age deviations of organ-specific model predictions or of individual protein levels are indicated on the horizontal axis. For each organ – lung, heart, brain, liver, kidney and pancreas – hazard ratios for the two best models (marine and purple) and two best normalized proteins (red and green) are shown.

Organ-specific models are typically developed by identifying organ-specific subsets of proteins secreted into blood plasma and trained to predict age or mortality risk (Goeminne et al., 2024). However, to the best of our knowledge, computational methods to select organ-specific proteins such as the “GTEx 4x” – assigning protein to organ/tissue if its expression is 4 times higher than in any other organ/tissue - method have never been rigorously evaluated in the context of organ-specific aging models, and, as a consequence, their ability to single out organ-specific proteins has been cast into doubt (Orsburn, 2024). We therefore systematically evaluated different methods for selecting organ-specific proteins using both transcriptome- and proteome-based datasets (Fig. 1B), see Methods. The GTEx 4x models in this paper are identical to those used in our previous work (Goeminne et al., 2024). The feature subsets derived from these methods vary in size and in the proteins assigned to different organs. However, across models based on these different protein subsets and trained on mortality and aging, several proteins are consistently selected by models as the largest contributors to the predictions. When these proteins’ model coefficients are multiplied by their average protein values to determine their contribution, the same proteins repeatedly emerge as top contributors (Supplementary Table 1, Supplementary Figure 1). For instance, heart models prioritize the NPPB protein, whereas lung models consistently include SCGB1A1, LAMP3, and SFTPD proteins (Fig. 1C). Furthermore, when assessing the stability of the models with 1000 iterations of LASSO on random data subsets, the same proteins are consistently included in the models in all iterations (Supplementary Table 2).

To assess the predictive capacity of proteins, we have used all proteins available in UK Biobank Olink panel. The hazard ratios of protein levels were compared to the aging models with highest hazard ratios for organ-specific diseases in six major organs: lung, heart, brain, liver, kidney and pancreas (Fig. 1F-K). The best organ-specific models for organs are predominantly mortality-based (with the exception of brain). However, when compared to individual proteins, the organ-specific models are less discriminative in predicting diseases. For example, the time-to-mortality heart model that based on 4-fold expression threshold calculated for limited organs (“GTEx 4x limited”) predicts heart failure with the highest hazard ratio (Supplementary Figure 2D) but predicts liver cirrhosis/fibrosis even better (Fig. 1G), similarly for lung-specific models (Fig. 1F). For brain, GFAP levels alone predict dementia with a hazard ratio of 1.70 versus 2.58 for the Feedforward Network (FFN) chronological brain aging model. But GFAP does not pick up signals of stroke, myocardial infarction, or cancer and has much weaker associations than the brain model with diseases manifesting in other organs (Fig. 1H). While for most organs, the best-performing proteins were a part of the respective organ-specific feature subsets for models, like NTproBNP for heart failure and LAMP3 for COPD, there are notable exceptions. For instance, ADM is the best predictor for both heart failure and kidney failure (Fig. 1G, 1J), but is expressed across multiple organs, like heart, lungs, kidney, liver, skin, bladder, etc. (GTEx Consortium, 2020).

Interestingly, cancer is not predicted well by any of the organ-specific models. However, when separated by specific cancer types, both organ-specific models and individual proteins can predict cancer. For example, KLK3 plasma levels predict malignant neoplasm of prostate with a hazard ratio of 3.70 (adjusted p-value < 1*10^-100^). Similarly, both LAMP3 alone and the lung-specific GTEx PMI AdaTiSS mortality-based model can predict upper lobe, bronchus or lung cancer with hazard ratios of 2.74 (adjusted p-value = 1.1*10^-40^) and 2.24 (adjusted p-value = 1.8*10^-62^), respectively (Supplementary Table 3). For the 24 types of cancer analyzed single proteins are always the best predictors, both in regard to hazard ratio and significance, with malignant neoplasm of kidney as the only exception. For this cancer type, the most significant predictor is a chronological model based on kidney-specific GTEx 4x stable features, even if the hazard ratio for protein HAVCR1 is still higher (2.53 vs 2.33, Supplementary Table 3).

In summary, single proteins can predict the hazard ratio of the analyzed diseases as well as or better than complex models, with the exception of brain, but the GFAP marker has an advantage over brain models when it comes to specificity.

### Single proteins predict multiple organ diseases and are robust to covariate adjustment

After establishing that single proteins predict diseases similar or better than organ-specific aging models, we investigated the specificity of these proteins and models as predictors of diseases from individual organs. The UK Biobank dataset includes over one thousand diseases annotated with ICD-10 codes and their first occurrence. We here solely focus on 373 diseases with over 100 occurrences post-sample collection. Some diseases affect multiple organs, while others primarily burden a specific organ. We here developed several variations (Fig. 1B) of our previously-reported GTEx 4x-based organ-specific aging models. We have shown before that such models simultaneously predict multiple diseases affecting an organ and prioritize organ-specific diseases (Goeminne et al., 2024). We now systematically grouped these 373 diseases by organs (Supplementary Table 4) and calculated the number of organ-specific diseases in the top 5% of hazard ratios for each model and protein (Fig. 2A). Organ aging models predict more diseases specific to that organ among the top 5% compared to conventional aging models (Supplementary Table 4). However, the proteins on which the models are based predict these diseases at a comparable rate (Fig. 2B-D, Supplementary Table 4). Notably, certain proteins, such as GFAP, which is brain-specific, predict more mental and nervous system diseases than the brain model, with the exception of the GTEx AdaTiSS model, which only retained GFAP itself as a predictor. Similarly, MSLN and SFTPD predict more lung-afflicting diseases among the top 5% of hazard ratios than all lung-specific aging models (Fig. 2C).

**Figure 2.**
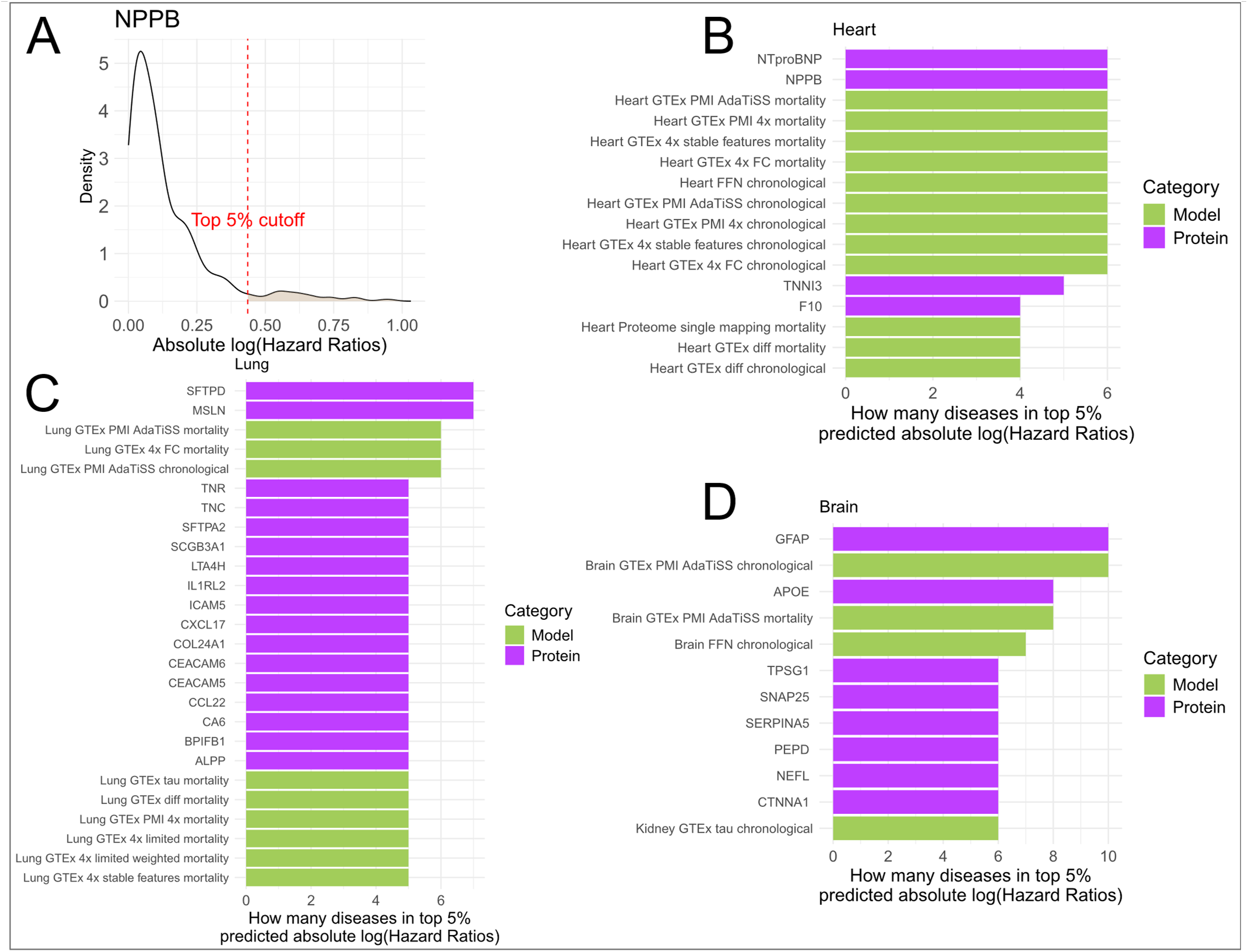
Single proteins generally predict more organ-specific diseases than protein-based organ-specific aging models. **A**. Distribution of absolute log(Hazard Ratios) for single predictor NPPB across 373 diseases that had at least 100 occurences after sample collection. The red line indicates the 5% cutoff of the diseases with the highest absolute log(Hazard Ratio). We calculate the number of heart-specific diseases that fall above the cutoff. The same analysis strategy was performed for all proteins and all aging models. **B-D**. Predictors with the highest number of organ-specific diseases in the top 5% of absolute log(Hazard Ratio) distribution in heart (**B**), lung (**C**) and brain (**D**). Number of diseases assigned to the categories: heart – 11, lung – 12, brain – 27. Colors indicate the type of predictor – aging model (green) or single protein (purple).

Prediction of diseases by individual proteins might be confounded by several factors. The Cox models in Fig. 2C – D account for sex, chronological age, and their interaction, but multiple additional factors can be considered when predicting diseases.

After accounting for BMI, waist circumference, smoking, drinking, the Townsend Deprivation Index, and assessment center, most diseases retain several top predictors, albeit with reduced hazard ratios and adjusted p-values (Fig. 3D-G). Similarly, when adjusting for 61 biochemical and hematological blood markers that are commonly used in clinical settings, such as cystatin C and HDL cholesterol, hemoglobin concentration and others, the number of significant interactions is further reduced, but multiple top predictors are retained (Fig. 3A-C). For instance, LAMP3 predicts COPD with an HR of 2.08 (adjusted p-value = 6.10*10^-245^) without additional covariate adjustment, an HR of 1.50 (adjusted p-value = 1.62*10^-55^) with adjustment for environmental factors, and an HR of 1.35 (adjusted p-value = 1.43*10^-11^) when adjusted for additional blood markers (Fig. 3G). NTproBNP is less confounded by covariates and predicts heart failure with an HR of 1.96 (adjusted p-value = 1.38*10^-273^), an HR of 1.88 (adjusted p-value = 3.70*10^-224^), and an HR of 1.83 (adjusted p-value = 2.92*10^-224^) with sex-age adjustment, additional environmental adjustment and additional adjustment for clinical blood markers, respectively (Fig. 3E).

**Figure 3.**
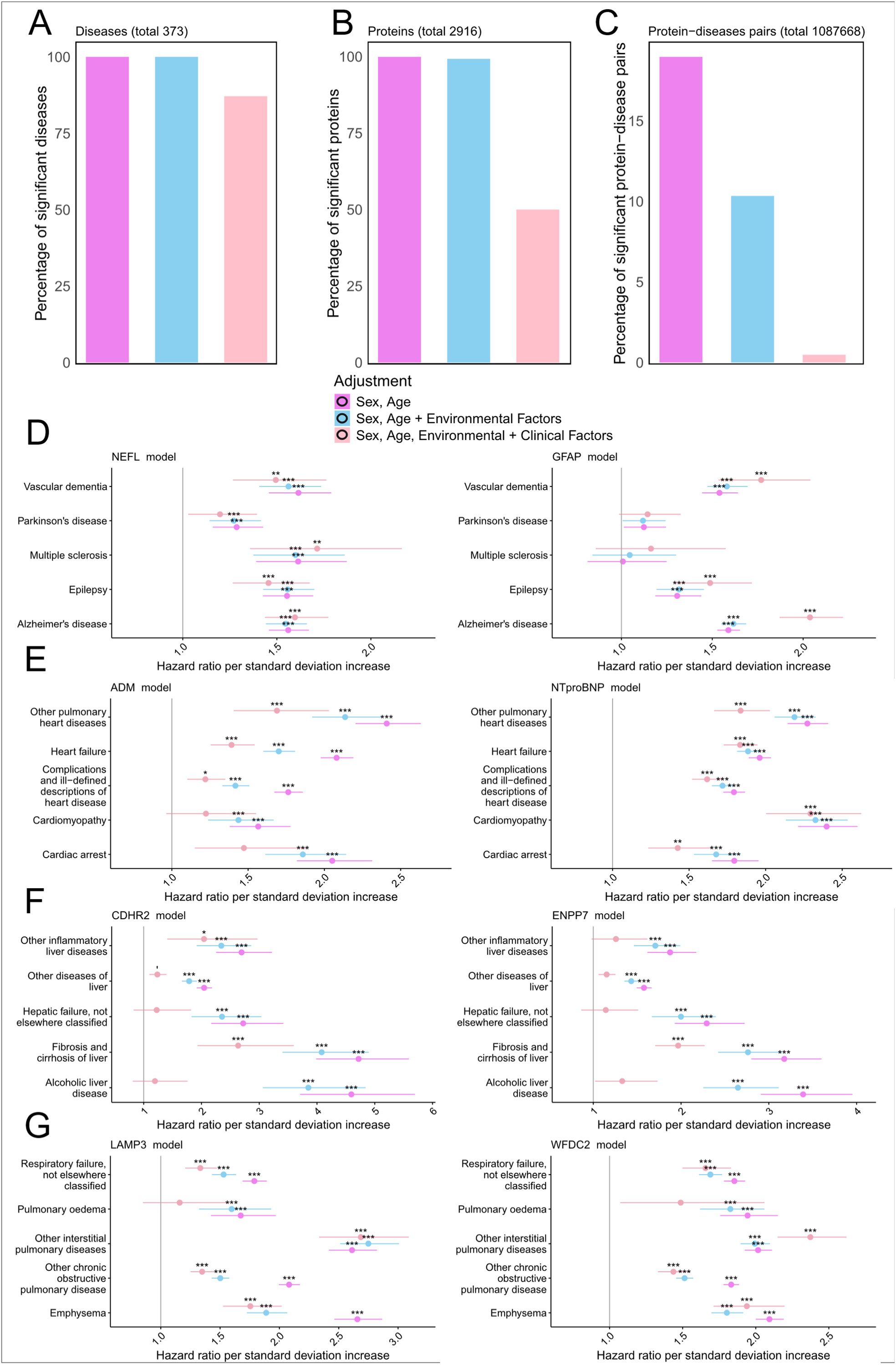
Selected best protein predictors for diseases are largely una#ected by covariate adjustment. A-C. Percentage of statistically significant (adjusted p-values < 5%) diseases (that have at least 1 significant protein association), proteins (at least 1 significant disease association) and protein-diseases interaction for each level of covariate adjustment: sex-age; sex-age and environmental; sex-age, environmental and blood markers (“clinical factors”). **D-G.** E@ects of covariate adjustment on the hazard ratios of the top-two single-protein predictors for the selected diseases in brain (**D**), heart (**E**), liver (**F**), and lung (**G**). Colors represent the adjustment type.

### Time-to-disease models outperform aging models, single protein markers and polygenic risk scores in disease prediction

We have demonstrated that individual biomarkers can often predict diseases more effectively than models trained solely to predict chronological age or mortality risk. However, models can also be specifically trained to predict diseases directly. We have trained Cox proportional hazards models with elastic net penalty on all available features (all proteins measured with Olink) to predict time to disease occurrence for a subset of age-related diseases: dementia, heart failure, kidney failure, COPD and Alzheimer’s disease. Figure 4A shows that time-to-dementia models directly outperform conventional and organ-specific aging models, as well as individual protein markers. The individual proteins NEFL and GFAP still outperform a conventional mortality-based model. However, a feed-forward neural model trained on the brain-specific proteins to predict chronological age performs comparably to the model that is trained to predict dementia on the same subset of proteins. Exploring different feature subsets of brain proteins (see Methods), we found that a FFN based on a small subset of 7 top-performing proteins (among those with highest coefficients in different brain aging models) - GFAP, NEFL, SEZ6L2, PODXL2, NPTXR, BCAN, and OMG – can predict dementia (Fig. 4D) and other brain-related diseases (Fig. 4E) similar to the neural network trained on 119 features, while being more selective for brain diseases (Fig. 4D) than other models.

**Figure 4.**
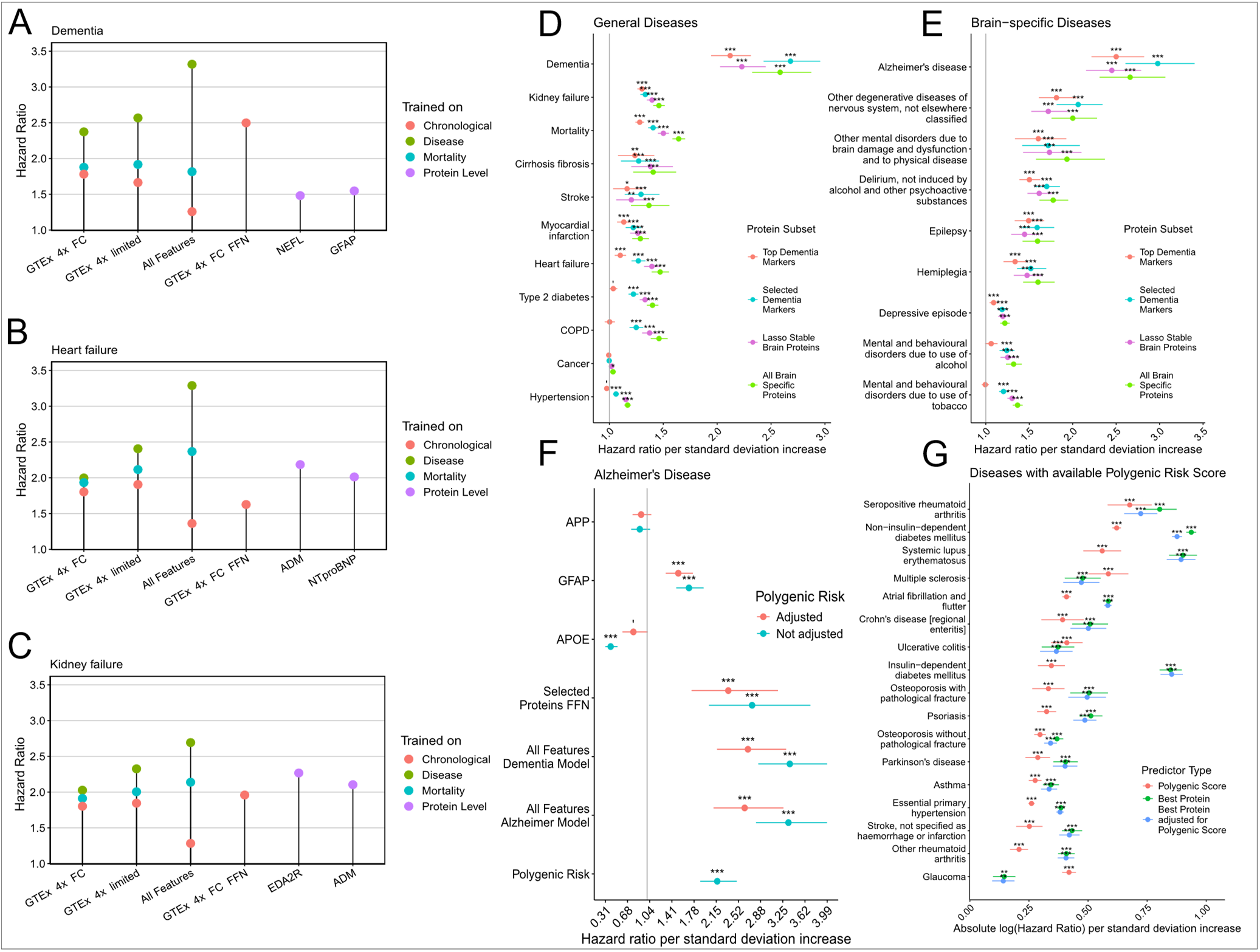
Brain chronological aging feed-forward neural networks perform comparably to models trained on diseases and can predict Alzheimer’s disease better than polygenic risk score. A-C. Hazard ratios per unit of standard deviation increase for predicting dementia, heart failure and kidney failure with models trained on chronological age, mortality, and disease outcome itself, as well as the best single-protein predictors. Dot colors represent the models’ targets, the horizontal axis shows the predictive method. GTEx 4x uses the GTEx 4x method for feature selection; GTEx 4x limited only considered the 8 largest organs for feature selection, “All features” models were trained on all features. These models are elastic net models; GTEx 4x FC FFN uses the 4x method with a FFN. The two remaining methods are the top two single-protein predictors. Feature selection for organ-specificity was done for brain (A), heart (B), and kidney (C). **D-E.** Comparison of different input feature subsets for the FFN brain model on hazard ratios in general diseases (**D**) and brain-specific diseases (**E**). The legend describes the protein feautre subsets used to train the neural network: top dementia markers – 5 proteins with the highest hazard ratios in predicting dementia individually, selected dementia markers – manually curated list of 7 proteins (GFAP, NEFL, SEZ6L2, PODXL2, NPTXR, BCAN, OMG), lasso stable brain proteins – proteins selected by LASSO in predicting chronological age on brain-specific proteins, as selected by the GTEx 4x method, in all 1000 runs, and all brain-specific proteins. **F.** Comparison of models and individual proteins in predicting Alzheimer’s disease. The vertical axis indicates the type of predictor – selected proteins, neural network trained on selected proteins and two elastic net models trained directly on dementia and Alzheimer’s disease, as well as the (Thompson et al., 2022) PRS reported in UK Biobank for Alzheimer’s disease. Color represents two types of Cox survival models –one with sex and age, and their interaction, as covariates (marine), and one with additional adjustment for polygenic risk (red). **G.** For diseases with available PRS in UK Biobank, we compare the predictive power of PRS (red), best protein (green) and protein adjusted for PRS (blue). The horizontal axis shows absolute log(hazard ratios) to allow direct comparisons of the strength of associations. It does not indicate the directionality between the predictor and the disease.

Similarly, training models directly on time to first occurrence of heart and kidney diseases results in the highest hazard ratios, with models including all features being predictably the best performers. Interestingly, the gap between hazard ratios of individual proteins and our disease models is much smaller for heart failure (Fig. 4B) and especially kidney failure (Fig. 4C), than for dementia (Fig. 4A).

Models trained on dementia and Alzheimer’s disease can predict Alzheimer’s disease with higher hazard ratio than individual proteins (Fig. 4F) like GFAP and APOE. However, APOE’s confidence interval is smaller, though its p-value is less than those of the disease models – 9.64*10^-13^ vs 3.00*10^-44^ for dementia model on the test subset of 8984 participants (restricted to one fold due to the computational intensity of predicting diseases using all features). The neural network on selected proteins is also capable of predicting Alzheimer’s diseases with HR = 2.74 (adj. p-value = 8.64*10^-11^), but with a wider confidence interval than other models. We then compared our predictions to Thompson et al., 2022’s Alzheimer’s disease polygenic risk score. This polygenic risk score (PRS) predicts the disease better than individual protein levels, but all multi-feature models demonstrate higher hazard ratios than the PRS. Adjusting for PRS lessens the prediction strength of all models and proteins, and in case of APOE significantly reduces the significance (Fig. 4F), as PRS already encapsulates a significant amount of genetic information associated with Alzheimer’s disease risk.

We extended this approach to other diseases. For the diseases with available PRS, we selected the best protein predictor (Supplementary Table 5) and calculated absolute log(hazard ratio) for the protein, PRS and protein adjusted for PRS (Fig. 4G). We found that most diseases show stronger associations with protein markers alone compared to polygenic risk (Fig. 4G). Glaucoma, ulcerative colitis and multiple sclerosis are the only diseases that are better predicted by PRS. The results of the likelihood ratio test for non-nested Cox models between the PRS and protein models (Supplementary Table 6) show that, for several diseases, the protein model provides a statistically significant better fit to the data compared to the PRS model: atrial fibrillation (p-value = 1.2*10^-14^), hypertension (p-value = 0.003), stroke (p-value = 0.002), psoriasis (p-value = 0.02), rheumatoid arthritis (p-value = 0.0005), lupus (p-value = 6.7*10^-5^), type II diabetes (p-value = 2.8*10^-14^), and type I diabetes (p-value = 5.0*10^-8^). When comparing the models with PRS only, and those including both PRS and best protein predictors, all tested diseases with available PRS information show significant better fit (p-value <5%) with the model that includes PRS and protein, indicating that protein capture additional information to PRS (Supplementary Table 6).

### Selecting stable disease predictors

While individual protein levels can predict multiple diseases, covariate adjustment can influence the ranking and significance of proteins for each individual disease. We sought to identify a panel of proteins that are always included among the top 10 predictors for each disease, irrelevant of the direction of the association, and find overlaps among our three covariate adjustment approaches, see section “Single proteins predict multiple organ diseases and are robust to covariate adjustment” of Results. 249 out of 373 diseases had at least one protein that was significantly associated with the disease in all three adjustment approaches. We only considered those consistent proteins for the panel that measure overall deterioration of heart, lung, brain, liver, kidney, pancreas, and/or overall mortality.

We further searched for specific protein predictors that are uniquely mapped to a single disease, or ICD-10 section. These disease-protein pairs were filtered by BH adjusted p-value below 0.05 and only proteins significantly associated with one disease were considered. After sex and age adjustment, there are 15 specific disease-protein pairs; for sex, age and environmental adjustment, there are 86, and for sex, age, environmental, and after additional adjustment for blood markers, there are 605 pairs (Supplementary Table 7). The only disease-protein pair that appeared in all adjustment lists is TMED1, which predicts gout with HR = 1.11 (adj. p-value = 0.002). When considering only adjustment for sex and age, as well as sex, age, and environmental factor adjustment, several more unique protein-disease pairs were identified. For example, diminishing levels of ACSL1 predict Crohn’s disease (HR = 0.64, adj. p-value = 0.002), an association that has previously been shown on the transcript level, (Knecht et al., 2016). Overall, specific protein-disease associations tend to have lower hazard ratios and higher p-values compared to top proteins that are associated with multiple diseases. For diseases grouped by ICD-10 diseases sections or chapters, the number of pairs increased only by a few disease-protein associations compared to considering all diseases individually (Supplementary Table 7).

Most proteins from the Olink panel are indeed associated with multiple diseases. Such diseases typically belong to different ICD-10 sections or even ICD-10 chapters (i.e. respiratory and circulatory). For example, ADM is significantly associated with 242 diseases (Fig. 5A) when adjusting only for sex and age. Out of the 242 diseases, the change in ADM levels affects the hazard for 238 diseases by at least 10% (HR >=1.1 or HR <=0.91), and for 90 diseases by more than 50%. Among the consistent proteins, i.e. significant in all models of adjustment, including just 6 proteins (when adjusted for sex and age) and 31 proteins (when adjusted for sex, age and environmental factors) is enough to significantly predict 300 diseases (Fig. 5A). Furthermore, when requiring a protein-disease pair to predict at least a 10% increase or decrease in disease hazard, 8 proteins are sufficient to predict at least 300 diseases when adjusted for sex and age, and 48 proteins are sufficient when adjusted for sex, age and environmental factors (Supplementary Figure 3A). A more stringent threshold of 50% risk increase/decrease shows that even when all proteins are considered, at most 177 diseases can be predicted significantly, but 8 proteins are enough to predict 100 diseases with this threshold (Supplementary Figure 3B and 3C). These proteins are ADM, CDHR2, CLEC7A, CD300LF, DPT, PILRB, and COL18A1.

**Figure 5.**
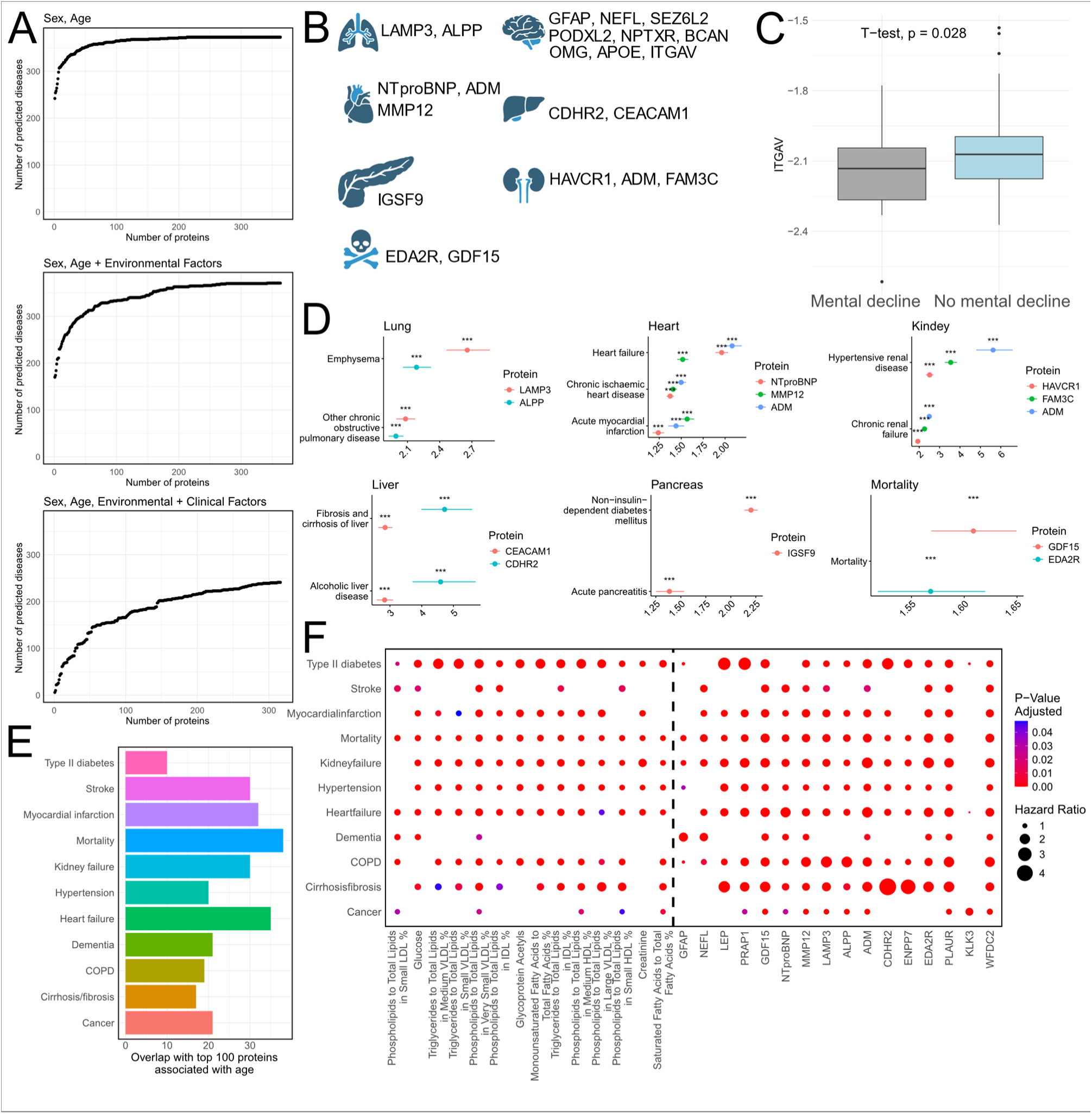
A small panel of 21 proteins can predict the health state of major organs. **A**. The number of significantly predicted diseases for different numbers of proteins from the consistent subset (i.e. the predictors of diseases that are robust to all covariate adjustments). Proteins are sorted by absolute log(hazard ratio), and each subplot represents adjustment type: sex-age, sex-age and environmental, sex-age and environmental and clinical blood markers. **B**. Proposed panel of 21 protein markers for predicting organ and body health state. **C**. Circulating ITGAV protein levels are significantly lower in individuals from PPMI cohort with and without cognitive decline onset. **D**. Unidirectional hazard ratios (expressed per unit of standard deviation) for the predictive markers from our 21-protein panel for their respective diseases associated with major organs and mortality, with the exception of brain, for which the previously shown FFN model is proposed. **E**. Overlap between top-100 predictive proteins for 10 age-related diseases and mortality, and top-100 proteins associated with chronological age. **F**. Comparison of metabolites’ and proteins’ predictive power for 10 age-related diseases and mortality. For each disease, top metabolite and protein predictors were selected. The black dotted line indicates the separation between metabolites and proteins.

As mentioned above, several previous studies explored the relationship between NMR metabolites and diseases (Gadd et al., 2024; Julkunen et al., 2023). The UK Biobank cohort includes 25,000 patients for whom NMR metabolites and Olink proteins levels were reported for the blood drawn at first visit. This allows for a direct comparison of metabolites’ and proteins’ predictive power. Figure 5F illustrates that individual metabolites have less predictive power than individual proteins. Among 350 diseases, 204 diseases had only proteins as top 10 significantly associated predictors. For some diseases, single metabolites come up as one of the best predictors, but they are always closely followed by single-protein predictors (Supplementary Table 9). For example, phospholipids to total lipids in large HDL percentage is the top predictor for menopausal and other perimenopausal disorders (HR = 1.16, adj. p-value = 0.00659), but is closely followed by protein predictor PAEP (HR = 1.15, adj. p-value = 0.0012).

The overall health state of the organism and organs can therefore be reflected with a low number of proteins. Based on these findings, we propose a platform of 21 proteins that can capture the health state of major organs, as well as general mortality risk (Fig. 5B, 5D, Supplementary Table 8). The panel includes broad predictors like ADM, EDA2R and GDF15, as well as organ-specific predictors like LAMP3 and NTproBNP. Our panel furthermore includes a set of proteins that serve as features for a neural network capable of predicting brain diseases comparably to models trained to predict dementia on all proteins (Fig. 4D). The brain set includes an additional protein: ITGAV, which is reported as a consistent predictor for Parkinson’s disease. We validated ITGAV in the external The Parkinson’s Progression Markers Initiative (PPMI) dataset, a comprehensive research project that collects longitudinal and multi-modal data on human healthy controls, individuals diagnosed with Parkinson’s disease, and prodromal patients. The PPMI also provides plasma Olink proteomics data, which facilitates direct validation of our protein biomarkers. Participants undergo regular assessments, and their cognitive status is recorded, categorized as normal, cognitive complaint, mild cognitive impairment, or dementia. Within the PPMI dataset, 73 patients with Olink data and recorded cognitive state were analyzed, 22 of whom experienced cognitive decline (Fig. 5C). ITGAV was found to predict cognitive decline with an HR of 0.56 (p-value = 0.03). These findings underscore the robustness of ITGAV as a biomarker for cognitive decline in Parkinson’s disease, thus validating its predictive power within the PPMI cohort. To measure the state of the lung, we selected LAMP3 and ALPP as the best predictors for COPD and other pulmonary disease (Fig. 1F), for heart, we selected ADM and NTproBNP as the best predictors of heart failure (Fig. 1G), and MMP12 as the best predictor of myocardial infarction (Supplementary Table 5). For kidney disease, HAVCR1, ADM, FAM3C together could significantly predict all kidney-related disease (Supplementary Table 8), as for liver, CDHR2 and CEACAM1 outperformed organ-specific aging models by a great margin, and stayed significant, even when accounting for alcohol consumption (Fig. 3F, Supplementary Table 5). For pancreas, we selected IGSF9 as a consistent and most significant predictor of diabetes (Supplementary Table 5), which is also predictive of acute pancreatitis (HR = 1.39, adj. p-value = 8.7*10^-9^). EDA2R and GDF15 are the strongest predictors of mortality (Supplementary Table 9). This proposed panel of 21 proteins (Fig. 5B) can significantly predict 336 diseases: 330 diseases with at least 10% change in hazard, and 137 diseases with >50% change in hazard per unit of standard deviation, when adjusted for sex and age, and their interaction (Supplementary Figure 4A). After adjustment for sex, age and environmental factors, there are 284 significantly predicted diseases, out of which 281 with at least 10% hazard change and 106 with >50% hazard change per unit of standard deviation (Supplementary Table 8, Suppl. Fig. 4B).

To showcase the performance of our panel, we compared its predictive power in mortality and diseases against other panels and the clinically-established biochemical and hematological blood markers (Fig. 6A-E). We then compared Cox elastic net time-to-mortality models trained on (1) our panel, (2) our panel including chronological age as a feature, and (3) the entire 2916 protein feature set available in UK Biobank to the PhenoAge mortality score, the metric which the epigenetic PhenoAge clock is trained to predict (Levine et al., 2018). We compared C-index, hazard ratios and specificity/sensitivity with the help of receiver-operating characteristic (ROC) curves in predicting mortality at a 10-year follow-up time point (Fig. 6A-C). C-index demonstrated best risk discrimination performance for the full protein set (0.80), followed closely by our 21-protein panel with a score of 0.78 (both with and without including age as an additional feature). PhenoAge mortality score, which also includes chronological age as feature, is comparable to the performance of the GDF15 single protein measurement (Fig 6A). Hazard ratios demonstrated similar trends with our 21-protein panel including chronological age slightly outperforming the full-protein model without chronological age (Fig. 6B). ROC curves show that sensitivity/specificity at a ten-year time point is comparable for these models, with only the PhenoAge score showing lower discriminative capabilities (Fig. 6C). To assess the predictive power of our panel for diseases to the alternative solutions, we compared it to the traditional clinical biochemistry and hematological blood panel, as well as to a recently published general and disease-specific mortality panel with 14 proteins (Sethi et al., 2023), referred also as Melamud et al. panel. We used the features in each of these panels to train Cox elastic net models towards the 10 diseases and mortality whose performances were compared on out-of-fold testing subsets. Protein panels outperformed the clinical blood panel for all diseases, both according to hazard ratios and C-index (Fig. 6D-E). With the exception of mortality, heart failure, stroke and myocardial infarction, where the 14- and 21-protein panels are tied, the 21-protein panel outperforms the 14-protein panel on all other diseases (Fig. 6D-E).

**Figure 6.**
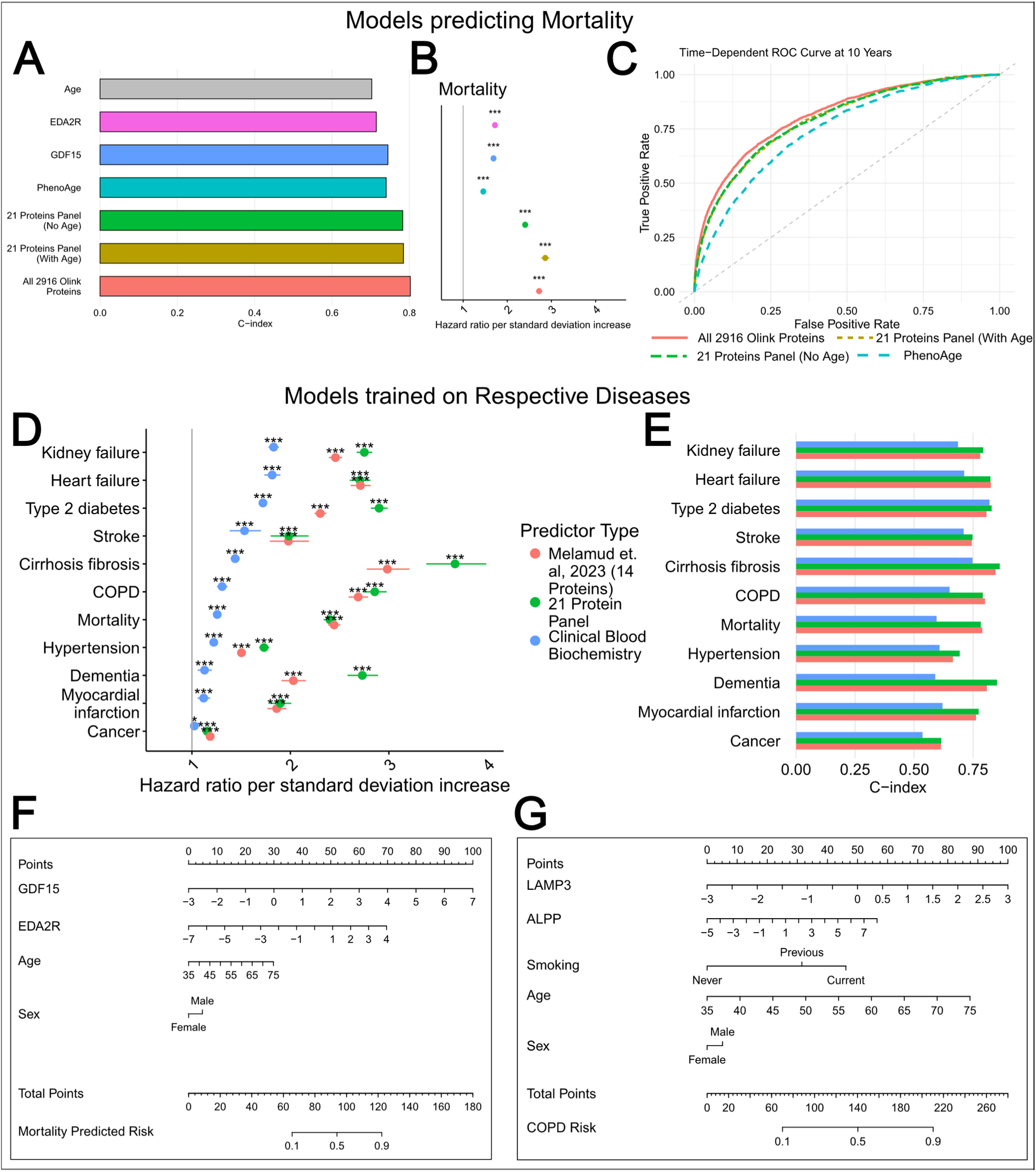
Comparison of the panel’s performance to other panels and established models. **A-B**. Comparison of C-index and hazard ratios in predicting mortality on test dataset by 21-protein panel trained on mortality (with and without age), full 2916 Olink panel trained on mortality, PhenoAge model, GFD15 and EDA2R levels, as well as chronological age alone. For every model (aside for chronological age), the hazard ratio for mortality corrected for age and sex is presented in figure **B.** The color legend is shared by **A** and **B. C.** Time-dependent ROC (receiver operating characteristic) curve for mortality after 10-year follow up for 4 models: all 2916 Olink proteins trained on mortality, 21-protein panel trained on mortality (with and without chronological age) and PhenoAge score. **D-E**. Hazard ratios and C-index of models trained on the respective diseases. Clinical blood markers, our 21-protein panel and the Melamud et al. 14-protein panel (Sethi et al., 2023) were trained on diseases and mortality and then compared on the test sets. The color legend is shared between **D** and **E.** All data in **A-E** is test data. **F-G**. Nomogram of mortality risk and COPD risk, respectively. For each value of a variable (e.g. Age or EDA2R), the points on top are assigned and summed to total points to generate a respective risk of mortality/COPD on the bottom.

While the panels can be used to train models to predict diseases, individual protein levels can be used to evaluate the risk of mortality or diseases like COPD, as represented in nomograms in Fig. 6F-G.

Finally, to assess the relationship between major diseases and aging, we calculated overlap between the 100 top predictors of these diseases and the 100 proteins most strongly associated with chronological aging (Fig. 5E). Each disease has a significant overlap (Supplementary Table 10) with chronological age markers, with heart failure leading with 35 overlapping proteins.

## Discussion

In this study we have utilized the proteomics dataset of the UK Biobank cohort of almost 50,000 subjects to show that individual protein levels can predict the disease state of organs better than organ-specific machine learning models trained on aging or mortality, and better than metabolites. In addition to the method of defining organ specificity by including proteins that are expressed in an organ at least 4-fold higher than in other organs (Uhlén et al., 2015), we applied a comprehensive set of organ-specific models, built on protein subsets, derived from different organ-specificity methods. We expanded the model space to include linear and non-linear methods, such as feed-forward neural networks (FFN). Additionally, we trained models directly to predict diseases. With the exception of the brain, major organs (heart, lungs, liver, kidney and pancreas) do not benefit from sophisticated models, and their deterioration can be predicted by a compact panel of 21 proteins.

Computational models to quantify biological age and disease risk have recently become a key focus in aging research, with growing interest in proteomics, particularly blood protein levels, which may offer an accuracy comparable to or better than epigenetic clocks (Ubaida-Mohien et al., 2023). Separate studies have focused either on chronological clocks, organ-specific clocks, disease clocks, individual proteins or metabolites, but there has been no effort to unite those findings (Argentieri et al., 2024; Carrasco-Zanini et al., 2024; Gadd et al., 2024; Goeminne et al., 2024; Julkunen et al., 2023; Oh et al., 2023; You et al., 2023).

Remarkably, expanding the pool of organ-specific proteins in models did not improve disease prediction (Fig. 1D-E, Supp. Fig.2A-D), as models consistently selected similar key proteins for each organ (Supp. Table 1, 2). For example, LAMP3 had the largest absolute contribution to lung-specific elastic net models (Fig. 1C), was selected as a lung-specific protein by almost all tissue-specificity methods, and consistently reappeared as top contributor in different models trained on those protein subsets (Supp. Table 1, Supp. Figure 1). By itself, it is capable of predicting COPD as well as models trained to predict mortality on 42 proteins (Fig. 1F). To the extent of our knowledge, LAMP3 has been investigated as a biomarker for an autoimmune disorder and a cell carcinoma, but not as a general predictor of lung diseases like COPD (Liao et al., 2015; Tanaka et al., 2020). In the case of heart diseases, NTproBNP, a known marker of cardiovascular disease (Welsh et al., 2022), outperforms most aging models, even if they include NTproBNP itself as a feature (Fig. 1G). As far as we know, a few selected proteins have not been explored in literature for their predictive capabilities of organ-associated diseases, like IGSF9 with respect to pancreatic diseases such as diabetes. CDHR2 was shown elevated with liver cirrhosis on the transcriptional level, but not as a plasma predictive marker (Chan et al., 2016), while FAM3C is a novel proposed urine biomarker for kidney disease in canine urine, but not in human plasma (González et al., 2023).

Since aging models are trained on chronological age or mortality, they prioritize the proteins that mostly correlate with age or mortality, which are not always the best predictors of disease, even when predicting organ-specific diseases with organ-specific aging models. In all organs, apart from the brain, mortality-based models outperform chronological models (Fig 1.F-K, Supp. Table 4), irrespective of the protein subsets used to train them. Specific diseases like heart failure, dementia, and chronic respiratory diseases contribute significantly to mortality in older adults, accounting for a large proportion of deaths (Centers for Disease Control C Prevention, 2022; Tinetti et al., 2012). This suggests that the presence and severity of these diseases are more directly linked to mortality than merely the age of the individual, and why mortality-based models are more suited for disease prediction. However, elastic net selects for correlation, not causality, and forces a linear relationship between predictors and the target even if it is absent in the real world (Mei et al., 2023). Features can have a negative interaction, concealing the signal of each individual contribution. Therefore, models could inadvertently conceal the predictive powers of individual proteins when it comes to organ-specific diseases.

Models trained directly on diseases outperform other types of models and individual protein predictors, but with varying magnitude (Fig. 4A-C). For some diseases and organs, like kidney failure, single proteins carry predictive power, comparable to models trained to predict the disease directly.

The major exception is brain aging. Not only are chronological aging models better at predicting brain-related diseases than mortality-based aging models, but it is also the only organ that benefits greatly from applying a neural-network model. It is consistent with Alzheimer’s diseases being only the seventh leading cause of death (Centers for Disease Control & Prevention, 2022). Indeed, the brain may integrate numerous organismal changes as a person ages, and they could be more closely tied to the passage of time (chronological age) than to mortality (Mattson & Arumugam, 2018). We have selected seven stable proteins that, when trained with feed-forward neural network on chronological age, are capable of predicting dementia and Alzheimer’s diseases almost as well as disease-specific models, trained on all proteins (Fig. 4D-E) and better than PRS (Fig. 4F). Several diseases, like stroke, lupus, diabetes, and Parkinson’s disease, can be predicted better by proteins than by the Tompson et al. PRS (Fig. 4G), and patients can benefit from tracking the protein levels in addition to relying on PRS. PRS estimate lifelong genetic predispositions, while protein levels are reflective of real-time physiological states, which could be more relevant for disease prediction.

Overall, individual proteins can more precisely separate organ-specific diseases from non-organ-specific diseases compared to organ-specific aging models (Fig. 2B-D, Supp. Table 4). For example, out of 373 diseases, GFAP predicts more brain-related diseases among its top 5% hazard ratios than brain-specific aging models, both chronological and mortality-based elastic net models, as well as FFN models trained on chronological age. GFAP is a known marker of brain-diseases, and while it lacks the capability to differentiate between the different diseases, it is sufficient to selectively measure brain deterioration (Mayer et al., 2013).

We have also shown that individual proteins remain predictive after adjusting for multiple covariates (Fig. 3), like BMI, drinking and smoking habits, biochemical and hematological blood biomarkers like cystatin C, glucose, HDL cholesterol and many others. This highlights the added value of protein measurements in predicting disease outcomes. Unlike traditional markers that we can already measure or observe, proteins provide complementary information that enhances risk stratification. This clinical relevance underscores the importance of incorporating proteomics into predictive models, as they capture aspects of disease risk not accounted for by existing covariates.

We assessed the specificity of disease predictions by single proteins and found that few proteins can be mapped univocally to a single disease (Supp. Table 7). Among proposed unique interactions are TMED1 and gout, an association that has not been reported in the literature, as well as ACSL1 and Crohn’s disease, whose role in the diseases of inflammatory bowel had been discussed before, though not at the plasma level and not in a predictive capacity (Knecht et al., 2016). Overall, due to the pleiotropic nature of many proteins, most are markers of multiple diseases across ICD-10 chapters, and a small number of proteins is sufficient to predict deteriorating organ states (Fig. 5A, Supp. Figure 3).

We therefore propose a compact panel of 21 Olink proteins capable of assessing aging of the major organs and general mortality (Fig. 5B). Our 21-protein panel strongly predicts mortality and disease outcomes, especially when benchmarked against other protein panels and traditional health indicators like the biochemical and hematological blood panel (Fig. 6A-E). When trained to predict mortality, the 21-protein panel, with and without age as a covariate, showed comparable performance to models trained on the full 2,916 protein feature set and outperformed the PhenoAge score (Levine et al., 2018), a widely used mortality predictor (Fig. 6A-C). Additionally, the 21-protein panel with age slightly outperforms the full-protein set without age, reinforcing the value of selected, targeted proteins when paired with key covariates (Fig. 6B). In disease prediction, the 21-protein panel consistently surpassed clinical blood panels in predicting various disease outcomes, as reflected by both hazard ratios and C-index (Fig. 6D-E). Although the panel showed similar performance to the 14-protein disease-specific mortality panel (Sethi et al., 2023) in diseases like heart failure, stroke, and myocardial infarction, it demonstrated superior predictive strength for other diseases (Fig. 6D-E).

Measuring the levels of this focused set of 21 proteins can be done at a fraction of the cost of a comprehensive 3,000 protein panel analysis, which makes testing more accessible and practical for clinicians. While the larger UK Biobank panel provides a more comprehensive view for researchers, our small 21-protein panel provides a comprehensive overview of an individual’s general health state.

We specifically propose an Olink panel, because there has been a reported discrepancy between different technique of measuring protein levels in blood (Eldjarn et al., 2023), and seeing as this study was based on single cohort and methodology, we cannot address generalizability of these findings to other platforms, like SomaLogic.

In conclusion, our study demonstrates that individual proteins, particularly a compact panel of 21 key proteins, can reliably predict organ deterioration and disease with a level of precision comparable to more complex, organ-specific aging models trained towards chronological age and mortality. These proteins offer a cost-effective and practical approach for assessing organ health, while also providing insight into disease risk across multiple organs. Our findings highlight the potential of plasma proteomics in disease prediction and underscore the need for further exploration of protein-based diagnostics in both clinical and research settings.

### Limitations of the study

A limitation of our study is the absence of validation of the proposed protein panel on an independent dataset. The prerequisites for validation of disease prediction are the availability of long-term disease information collection of the participants after Olink sample collection, which complicates the search for validation datasets. Utilizing only UK Biobank and only a single proteomics platform is an additional limitation. It would be important in the future to validate proteins in our models with other methods and in additional ethnicities.

## Methods

### Human cohorts

UK Biobank is a large-scale biomedical resource containing genetic, health, and lifestyle data from 500,000 UK participants, aged 40-69, collected between 2006-2010 (Sudlow et al., 2015). UK Biobank stores longitudinal data about participants’ health adverse events and mortality. For 54,265 participants, the proteomic profiles of the blood plasma samples collected at the admission were characterized by the Pharma Proteomics Project (Sun et al., 2023). The protein levels were measured with the Olink technology - Proximity Extension Assay and provided by UK Biobank in Normalized Protein eXpression (NPX) values. The sample collection, Olink methodology, data quality control and processing have been described in detail in Supplementary Information of (Sun et al., 2023). We utilize the provided values without further processing, unless stated otherwise.

The Parkinson’s Progression Markers Initiative (PPMI) is a multi-center, observational study that tracks the progression of Parkinson’s disease across its various stages (Marek et al., 2011). The PPMI (Tier 1) provides plasma Olink data, facilitating the direct validation of protein biomarkers. The PPMI database includes healthy controls, individuals diagnosed with Parkinson’s disease, and prodromal patients. Participants undergo regular assessments, and their cognitive status is recorded, categorized as normal, cognitive complaint, mild cognitive impairment, or dementia. The publicly available data (Tier 1) provides Olink measurements of plasma protein levels in samples collected at different timepoints for participants. The dataset was used to validate a link between Parkinson’s disease progression and identified disease marker.

### Data processing

#### UK Biobank

The protein levels were available for 53,104 participants and 2,923 proteins at the date of access (19.04.2024) for the blood plasma sample collected at the base level (instance 0). The participants were excluded if they had missing data in more than half of the proteins, and proteins were excluded if they had more than 10% missing data points across the dataset. The filtering resulted in subset of 44,952 participants and 2,916 proteins. The age was determined for the date of sample collection. Missing values were imputed with k-Nearest-Neighbors imputation algorithm (k=10). For model training, the values were imputed after separation into training and test for each of the 5 folds.

Status of death was determined from UK Biobank field 40000, the censoring date for participants without reported death date was the latest death date reported in UK Biobank. Time to death or censoring was determined by subtracting the sample collection date (field 53) from the death date or censoring date, respectively. The UK Biobank provides dates for first occurrences of 1,165 health-related outcomes (Category 1712). We excluded outcomes related to pregnancy, perinatal period and congenital malformations. We further filtered the diseases to include only those with at least 100 instances after sample collection date (field 53), resulting in 373 diseases. We derived status and time-to-event the same way as with mortality, with an additional step: if the participant had died before the latest occurrence of the disease, the censoring date was set to the death date.

For our subset of 10 common age-related diseases, we assessed first occurrence of heart failure (ICD-10 code - I50), stroke (I64), type II diabetes (E11), chronic obstructive pulmonary disease – COPD (J44), fibrosis/cirrhosis of liver (K74). Additionally, the following diseases were grouped by ICD-10 codes: dementia (F00-F03), kidney failure (N17-N19), hypertension (I10, I15), myocardial infarction (I21, I22), and any cancer (UK Biobank field 40005).

#### PPMI

We used baseline Olink measurements provided for 73 participants with available cognitive state, which includes 22 participants with recorded cognitive decline (cognitive complaint, dementia or mild cognitive impairment) after sample collection. The Olink data was accessed on 17.04.2024, cognitive status data on 18.06.24. Missing protein levels were imputed with the k-Nearest-Neighbors imputation algorithm (k=10).

### Organ-specific subset of proteins

Previously, we and others have applied the GTEx 4x approach, where we defined organ-specific proteins as those proteins for which the GTEx transcript expression in an organ is at least four times higher in that organ as compared to all other organs (Goeminne et al., 2024; Oh et al., 2023, 2024), whereby GTEx tissues were mapped to organs based on Supp. Table 2 from Oh et al. (2023). For this study, we extended this approach by not only considering tissue-level transcriptomics (GTEx Consortium, 2020), but also tissue-level proteomics (Jiang et al., 2020; Prakash et al., 2022) and single-cell transcriptomics (Consortium* et al., 2022), and including two additional specificity methods besides 4x: Tau (Lüleci & Ylmaz, 2022; Yanai et al., 2005) and AdaTiSS (Wang et al., 2021).

We investigated eleven methods of defining tissue-specific protein subsets and their impact on disease prediction hazards. The 4x fold expression method on GTEx transcriptomic data (GTEx 4x FC) served as a baseline for comparison. For this method we have used three adaptations: (1) limiting organs to only include the eight largest ones – brain, heart, liver, lung, kidney, pancreas, spleen and skin - (GTEx 4x limited), (2) adjusting for their organ weight (GTEx 4x limited weighted), and (3) the stable subset for organ-specific proteins is defined on 1000 iterations of LASSO (GTEx 4x stable features) – only the proteins that were selected by all 1000 iterations of predicting age based on the GTEx 4x subset were included in the subset of GTEx 4x stable features. Other methods to define protein specificity include tau (GTEx tau) and AdaTiSS (GTEx pmi AdaTiSS) metrics, as well as differential analysis (GTEx diff) – proteins whose expression was significantly 1.5 higher than in other tissues based on GTEx data. We also included two approaches where we first normalized the GTEx data for post-mortem interval (PMI) (GTEx pmi 4x, GTEx pmi AdaTiSS). Aside from the GTEx transcriptomics data, we used tissue-specific mass-spectrometry-based proteomics data: AdaTiSS method applied to post-mortem interval adjusted data based on the human tissue proteome map – Proteome AdaTiSS (Jiang et al., 2020), as well as proteins that were mapped by the authors to a single organ by integrative analysis of proteomics datasets – Proteome single mapping (Prakash et al., 2022). We also used the endothelial subset of the Tabula Sapiens transcriptomics dataset, where gene transcripts were mapped to different organs based on differential analysis in the endothelial cell type subset - Endothelial differential (Consortium* et al., 2022).

The Tau statistic, proposed as one of the more robust methods for tissue specificity by Kryuchkova-Mostacci C Robinson-Rechavi, 2017, is calculated as follows:

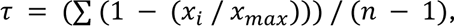

where x_i_ is the expression level of the gene in tissue i, x_max_ is the maximum expression level of the gene across all tissues, and n is the number of tissues being compared. A value close to 1 indicates high tissue specificity, meaning the gene is expressed predominantly in one organ. We have utilized the extended Tau method, that provides degree of specificity to each organ, (Lüleci & Ylmaz, 2022) to assign organ-specificity based on the GTEx transcriptomics data - GTEx Tau. The threshold for specificity is per default 0.85.

The AdaTiSS method is based on a Gaussian mixture model. It accounts for heterogeneities in gene expression or protein abundance and addresses the presence of outliers. The method applies a data-adaptive estimation procedure that uses density-power-weighting to estimate population parameters under unknown outlier distributions and non-negligible outlier proportions (Wang et al., 2021). We used AdaTiSS for GTEx pmi AdaTiSS, for Proteome AdaTiSS, we used the protein-to-tissue mapping by Jiang et al., 2020, that we assigned to organs. The mapping of proteins to organs with all 11 methods is available in Supplementary Table 11.

Additional subsets for brain-specific FFN models were created based on the results of Cox hazard models predicting dementia using individual proteins. Proteins with the strongest contributions to dementia prediction, identified from brain-specific and conventional feature sets, were further analyzed to assess how their inclusion and interaction parameters influenced Cox model predictions. Subsets containing 5 to 10 proteins were tested as feature sets for FFN chronological models to evaluate their predictive power for dementia and Alzheimer’s disease. The optimal feature set, consisting of 7 proteins – “Selected Proteins FFN”, was selected based on performance.

### Models

The organ-specific and conventional models (all proteins) were trained to predict chronological age and time-to mortality. The data for 44,952 participants was separated randomly into 5 folds. The train and test datasets were imputed with the k-nearest neighbor algorithm (k = 10) after separation. All reported results of models evaluation in the study are for test subsets. Reported Pearson correlation is a median across test subsets. The residuals for hazard analysis are calculated for the five combined test subsets using the models trained on their corresponding training sets (out-of-fold approach).

For models predicting chronological age, we used three related linear regression-based methods: Least Absolute Shrinkage and Selection Operator (LASSO), ridge, and elastic net (i.e. a combination of the LASSO and ridge). We utilized grid searching to optimize the L1 ratio parameter and the “alpha” parameter, a multiplier of the L1 and L2 penalty terms. Additionally, we tested feed-forward neural networks (FFN) for predicting chronological age. More specifically, we used a sequential model with 3 dense layers and tested following optimizations: rectified linear unit function (“relu”), the hyperbolic tangent function (“tanh”) and the logistic sigmoid function (“sigmoid”). Our FFN model also included one final prediction dense layer with linear activation. The optimizer was Adam with tested learning rates 0.01, 0.001, and 0.0001, batch size 32, and epochs 100.

For mortality and disease risk prediction, we used Cox’s proportional hazard’s models with elastic net penalty. The optimization parameters are the L1 ration and alpha parameters, as for the chronological models. All models were optimized to minimize the mean absolute error. When predicting diseases on all proteins, we only used a single fold due to the greatly increased computational time. The results are reported for the test subset.

### Hazard analysis

We utilized Cox Proportional-Hazards models to assess the effect of the predictors on the time to the first occurrence of a disease. For individual metabolites and protein levels, we normalized the NPX values by their standard deviations, for organ-specific models we used normalized residuals of prediction values regressed on chronological age (“age deviations”). Protein and metabolite values were adjusted for age in all Cox models, also removing the differences in levels occurring with age.

For each disease, we analyzed all available predictors – all chronological and mortality models on 11 feature subsets and FFN chronological models based on GTEx 4x for the organs available in each feature subset, proteins and NMR metabolic biomarkers (UK Biobank category 220).

Additionally, for 10 major age-related diseases and mortality, we trained the models on hematological (UK Biobank category 100081) and biochemical (UK Biobank category 17518) blood panel measurements, our panel of 21 proteins and a 14-proteins mortality panel (Sethi et al., 2023) to predict directly those 10 diseases and mortality.

Proteins and models’ residuals after regressing on chronological age were recorded for the full set of 44,952 participants, and are termed age deviations throughout the manuscript, in line with the recommendations of (Moqri et al., 2023). NMR Metabolites were reported for a subset of 24,151 participants, therefore comparison between hazard ratios of proteins and metabolites are performed only on this subset.

All our Cox proportional-hazard models are specified as follows:ℎ(***B***) = ℎ_0_(***B***) ∗ exp (***βX***)

Herein, ***B*** represents an *n*\*1 column vector of survival times *t*_*i*_, with *i* a 1: *n* index unique for every subject and *n* the number of subjects. ℎ(***B***) is the hazard function, which, depending on the outcome of interest, represents the risk of dying or the risk of being diagnosed with a specific disease at time *t*. ℎ_0_(***B***) contains the baseline hazards, i.e. the hazards if all covariates are 0. exp is the exponential function. X is an *n* * *p* matrix containing the *p* covariates for all *n* subjects. ***β*** is a *p* *1 column vector containing the *p* effect sizes corresponding to every covariate. Covariates always include a fixed intercept, and, unless specified otherwise, include a continuous term for chronological age, a dummy variable denoting sex, and an interaction term between age and sex. Besides these, we also performed analyses in which environmental factors where included as covariates as well, and analyses in which both environmental factors and biochemical and hematological blood markers were included as additional covariates.

All UK Biobank fields included for the environmental and the “clinical factors” - biochemical and hematological blood marker covariates are reported in Supplementary Table 11. For the analysis of polygenic risk, we calculated the hazard risk for polygenic risk scores adjusted for sex, age and interaction, and the hazard risk for protein levels with polygenic risk as an additional covariate to age, sex and their interaction. We report hazard ratios or absolute log(hazard ratios) per one unit of standard deviation of the predictor of interest. All p-values were adjusted for multiple testing with the Benjamini-Hochberg false discovery rate (FDR) method, and we always controlled the Benjamini-Hochberg FDR at the 5% level.

### Protein contributions

To evaluate the contribution of individual proteins in models to predicted values, we multiplied linear model coefficients by average protein values in case of linear models and in case of neural networks, we utilized the Python SHAP module (version 0.46.0) to explain the output of the neural networks. SHAP (SHapley Additive exPlanations) is a method used to interpret the output of complex machine learning models by assigning each feature an importance value for a particular prediction (Lundberg & Lee, 2017).

### Predictors’ preference for organ-specific diseases

For the analysis of preference of organ-disease predictions by models and proteins, we looked at the highest absolute log(Hazard Ratios) for diseases for each predictor – protein or a model. For lung, heart, brain, kidney, liver and pancreas, we manually grouped the diseases by organs (Supplementary Table 4), and calculated the number of organ-specific diseases in the top 5% of associations, sorted by absolute log(hazard ratio).

### Software

Most of the analysis was conducted in R (4.3.3 version, 29.02.2024) on 64-bit Windows 10 System. Cox Proportional-Hazards model were fit using the R survival package (3.5-8 version). All machine learning models were trained in Python 3.12.3, using the scikit-learn (1.3.2), and keras (3.3.3) modules.

## Acknowledgements

This study is supported by the National Institute on Aging and Hevolution Foundation. The research has been conducted using the UK Biobank Resource (application number 21988).

