## Supplementary Figures 1-4 for "A compact protein panel for organ-specific age and chronic disease prediction"

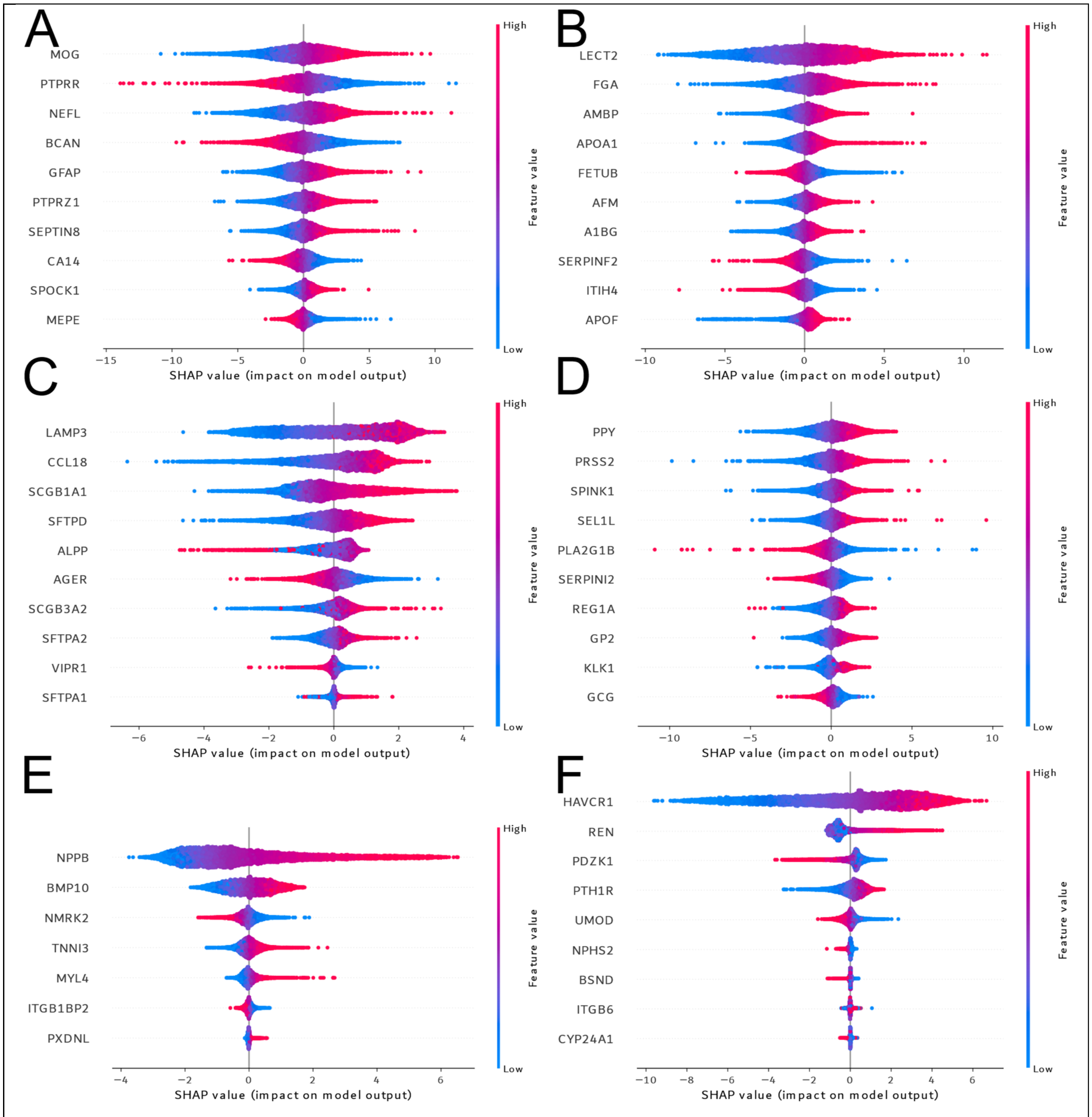


**Supplementary Figure 1. Protein feature importance in organ-specific feed-forward neural networks.**

**A-F.** For brain, liver, lung, pancreas, heart and kidney respectively, we applied SHAP to investigate the contribution of proteins to chronological age predictions by feed-forward neural networks. The method was applied to models trained on the first fold and test data not included in the training. The proteins are sorted by their importance. X-axis represents the impact on the model for each sample. Each dot represents individual samples, and the color indicates their relative value. Only the ten most important proteins are displayed per organ model.

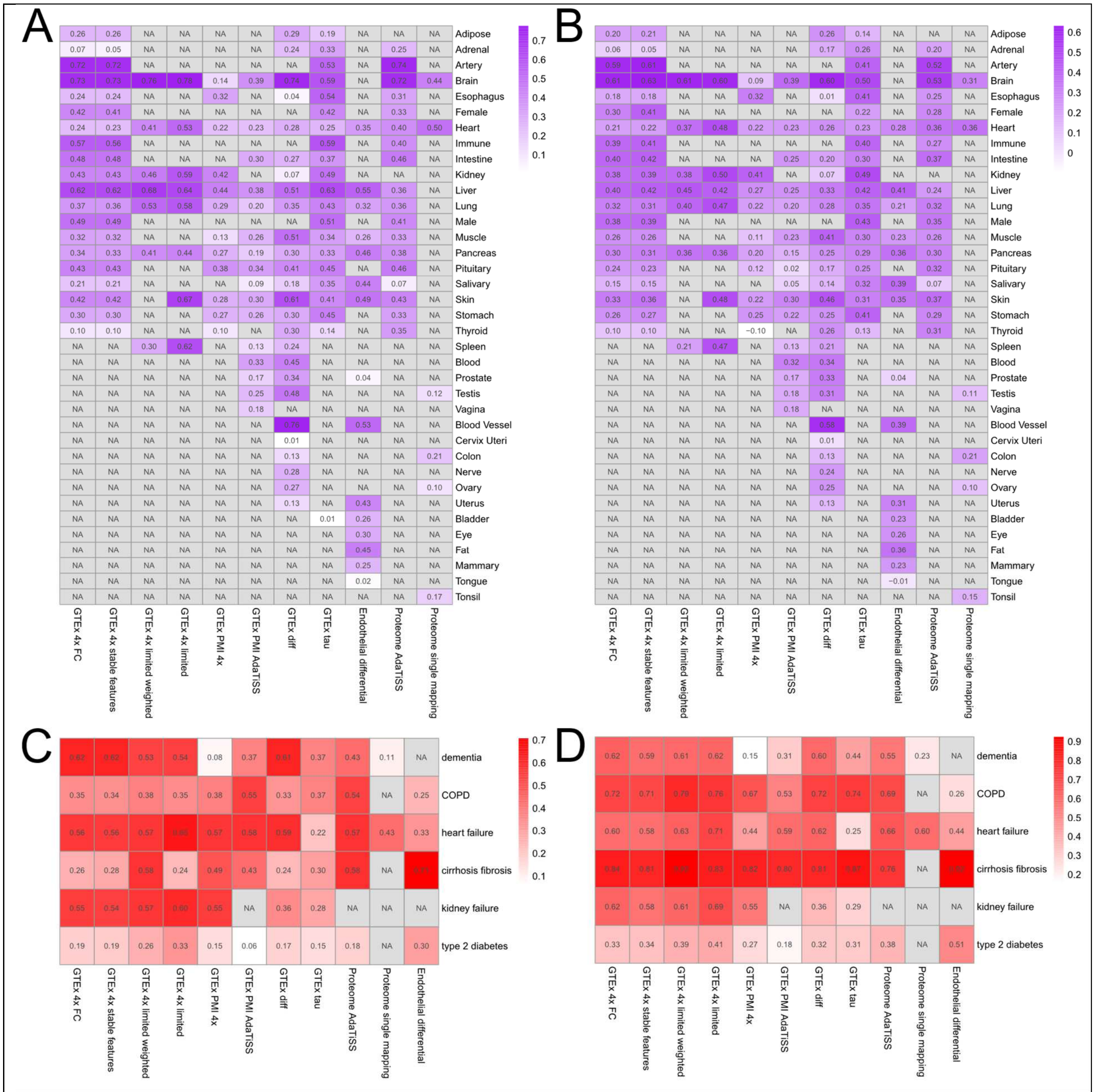


**Supplementary Figure 2. Comparison of organ-specific chronological and mortality models' performance.**

**A-B.** Pearson correlation with chronological age for chronological and mortality models respectively. The x-axis represents the type of subset approach for selecting proteins for respective organs in y-axis. The organs are not present in all models due to the difference in organ mapping and availability of tissues in different datasets, as well as presence of proteins that passed the threshold of "specific proteins" for the organ for the given method. **C-D.** Absolute log(hazard ratio) for selected organs in predicting diseases based on chronological or mortality residuals respectively. Organ to disease mapping: dementia-brain models, COPD – lung models, heart failure – heart models, cirrhosis/fibrosis – liver models, kidney failure – kidney models, type 2 diabetes – pancreas models.

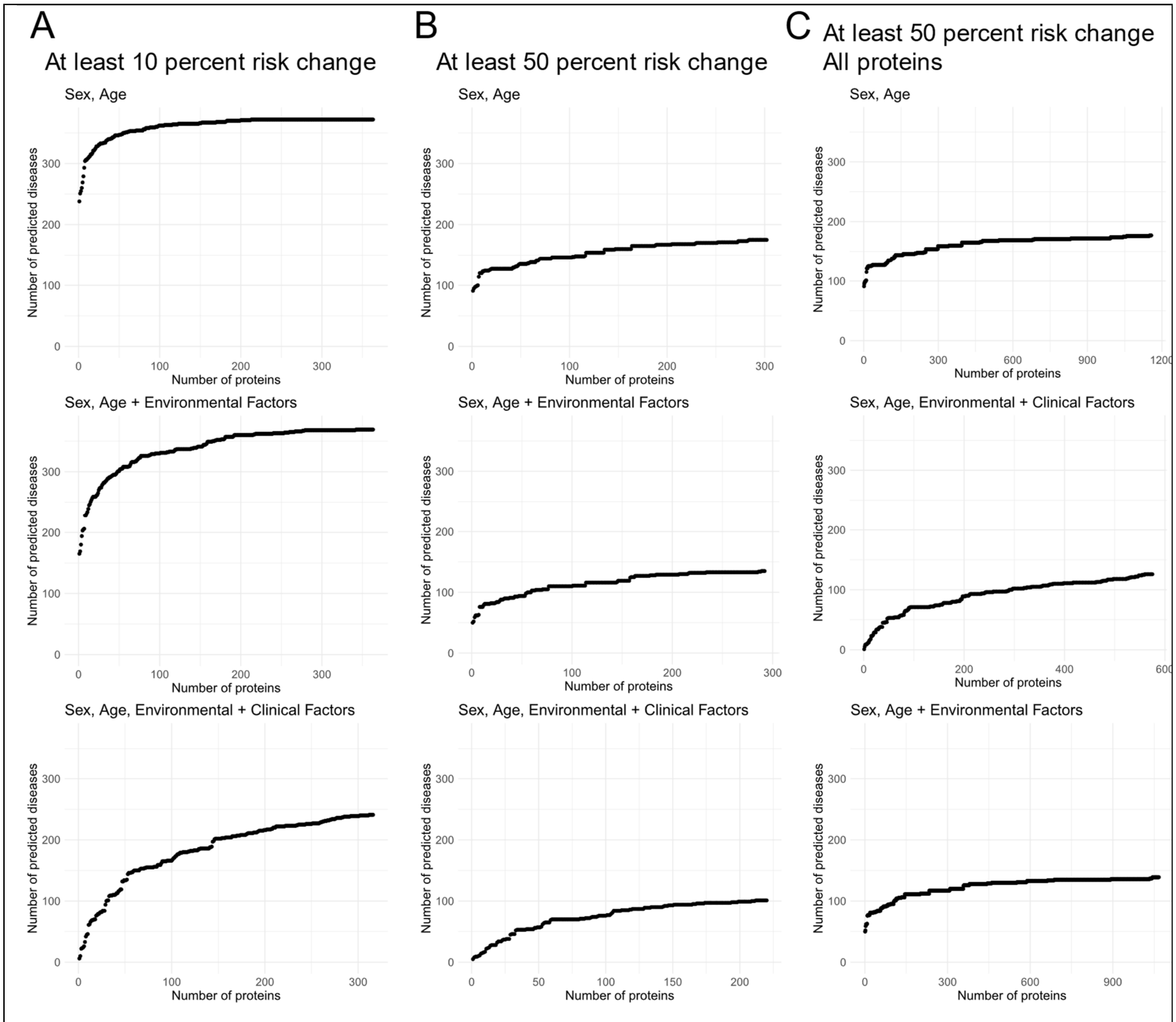


**Supplementary Figure 3. Number of proteins significantly associated with diseases.**
**A-C.** Proteins sorted by absolute log(Hazard Ratio). The number of diseases predicted by adding additional proteins is shown on y-axis. Only the disease associations that increase/decrease diseases risk by 10% (**A**) or 50% (**B, C**) are considered. **A** and **B** include only consistent protein predictors (i.e. predict at least one disease in all adjustment models), **C** includes all available proteins.

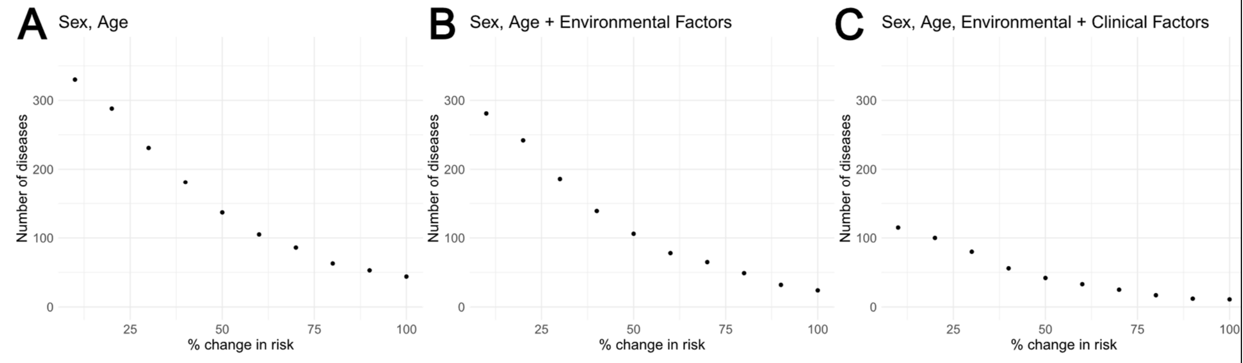


**Supplementary Figure 4. Number of diseases significantly predicted by 21-protein panel with the respective change in risk.**

**A-C.** Diseases significantly predicted by a 21-protein panel are filtered based on the change in the risk (on x-axis) and the number of these diseases for each change in risk threshold is displayed (y-axis). This is repeated for three types of adjustment: sex-age (**A**), sex-age and environmental factors (**B**) and sex-age, environmental and circulating clinical factors (**C**).
